# Distinguishing response from stimulus driven history biases

**DOI:** 10.1101/2023.01.11.523637

**Authors:** Timothy C. Sheehan, John T. Serences

## Abstract

Perception is shaped by past experience, both cumulative and contextual. Serial dependence reflects a contextual attractive bias to perceive or report the world as more stable than it truly is. As serial dependence has often been examined in continuous report or change detection tasks, it unclear whether attraction is towards the identity of the previous stimulus feature, or rather to the *response* made to indicate the *perceived* stimulus value on the previous trial. The physical and reported identities can be highly correlated depending on properties of the stimulus and task design. However, they are distinct values and dissociating them is important because it can reveal information about the role of sensory and non-sensory contributions to attractive biases. These alternative possibilities can be challenging to disentangle because 1) stimulus values and responses are typically strongly correlated and 2) measuring response biases using standard techniques can be confounded by *context-independent* biases such as *cardinal bias* for orientation (i.e., higher precision, but repelled, responses from vertical and horizontal orientations). Here we explore the issues and confounds related to measuring response biases using simulations. Under a range of conditions, we find that response-induced biases can be reliably distinguished from stimulus-induced biases and from confounds introduced by *context-independent* biases. We then applied these approaches to a delayed report dataset (N=18) and found evidence for response over a stimulus driven history bias. This work demonstrates that stimulus and response driven history biases can be reliably dissociated and provides code to implement these analysis procedures.

## Introduction

Perceptual reports can be shaped by past stimuli and actions - the visual system exploits this information to support efficient information processing. To this end, the visual system expends less energy processing expected stimuli and can rely on priors to facilitate processing of new sensory information (Mumford 1994; Oliver 1952; Olshausen and Field 1996). However, even though these adaptive mechanisms support more efficient processing on average, they also lead to a collection of perceptual biases.

For example, over developmental or evolutionary time scales perceptual processing has adapted to represent frequently encountered stimulus features such as vertical and horizontal orientations with greater precision than off-cardinal oblique orientations (the *oblique effect*). While this resource allocation supports more efficient processing in early visual cortex, it also gives rise to a phenomenon of *cardinal bias* where perceptual reports are repelled from vertical and horizontal orientations (Girshick, Landy, and Simoncelli 2011; Wei and Stocker 2015). Importantly, *cardinal bias*, as well as the *oblique effect*, are thought to be based on long-term exposure to natural image statistics and are highly stable across time (Henderson and Serences 2021). Hence, we use the term *context-independent biases* to refer to this and related phenomena.

In addition to these *context-independent* biases, dynamic perceptual biases can also emerge based on exposure to recent stimulus features. For instance, viewing a stable image feature for an extended period can lead to a suppressed neural response to that feature (Dragoi, Sharma, and Sur 2000; Kohn and Movshon 2004; Patterson, Wissig, and Kohn 2013). Given that stimuli are generally stable across time, these *adaptation* effects are also thought to contribute to *efficient coding* as fewer neural resources (i.e., spikes) are dedicated to processing expected stimulus features (Barlow 1961; Benucci, Saleem, and Carandini 2013; Felsen, Touryan, and Dan 2005). However, attenuated responses in neurons tuned to the viewed stimulus can bias neural population response profiles away from the adapting stimulus. This neural repulsion is the likely source of perceptual repulsion effects seen in well-known phenomena such as the waterfall illusion or the tilt after-effect (Anstis, Verstraten, and Mather 1998; He and MacLeod 2001).

Interestingly, and in contrast to typical adaptation-induced repulsive biases, the repetition of similar stimuli can sometimes lead to an attractive or assimilative bias known as hysteresis or serial dependence (Chopin and Mamassian 2012; Cicchini, Anobile, and Burr 2014; Corbett, Fischer, and Whitney 2011; Fischer and Whitney 2014). Typically, attractive serial dependence emerges with briefly presented or near-threshold stimuli that are hard to perceive, as opposed to longer exposure to high contrast stimuli that usually leads to adaptation and perceptual repulsion (Chopin and Mamassian 2012; Cicchini, Mikellidou, and Burr 2017; Fritsche, Mostert, and de Lange 2017; Maus et al. 2013). These attractive biases can be explained by invoking a Bayesian prior for stimulus stability over short time scales (van Bergen and Jehee 2019; Cicchini and Burr 2018; Fritsche, Spaak, and de Lange 2020; Pascucci et al. 2019). Given this prior for environmental stability, the precision of near-threshold stimuli can be improved by biasing reports towards recently viewed features (Cicchini and Burr 2018; Fritsche et al. 2020; Sheehan and Serences 2022). However, even though attractive biases are observed across a host of stimulus/task domains, their ultimate source is still debated.

Here we address a set of key unanswered questions related to efficient information processing in the human visual system. First, do attractive serial dependence effects depend on the physical identity of recently seen features, or on the responses made to report the identity of recently seen stimuli? Second, how do attractive serial dependence effects interact with adaptation and *context-independent* factors like *cardinal bias*? Parceling out sensory and motor contributions from these other perceptual biases is critical to better understanding the source of the effect because these factors all jointly contribute to measured perceptual reports.

Disentangling sensory from motor/decisional contributions to attractive serial biases is particularly challenging because most studies of serial dependence have employed delayed recall paradigms where responses are highly correlated with the presented stimulus feature. For example, in a typical task a participant is instructed to report the orientation of a remembered orientation using a mouse pointer. Their response will ultimately be driven by the integration of sensory evidence on that trial, adaptation induced by previous stimuli, *context-independent* biases (e.g., *cardinal bias*), and random errors accumulating from other unmeasured sources.

These will cause the response to deviate from the stimulus orientation but only by a few degrees such that even for a low performing participant, stimulus identity and the associated responses will still be highly correlated (r_circ_=0.63, σ=21.4° for an example continuous report dataset which we analyze in more detail below).

Most studies of serial dependance have focused only on the influence of the previous stimulus and claim that it is the processing or perception of the physical stimulus that induces attractive biases (Cicchini and Burr 2018; Cicchini et al. 2017; Fischer and Whitney 2014; Manassi et al. 2018). However, the emerging consensus is not so straightforward. One recent study found evidence that responses are simultaneously repelled (due to adaptation) and attracted (due to the application of Bayesian priors) to past stimuli but at different timescales, leading to both attractive and repulsive effects (Fritsche et al. 2020). In contrast, other work suggests that it is the previous decision, not the stimulus per se, that leads to attractive serial biases (Pascucci et al. 2019).This finding is consistent with subsequent studies that have simultaneously modeled the influence of both the previous response *and* the previous stimulus and found that reports are simultaneously attracted to previous responses and repelled from previous stimuli, providing an extra layer of distinction between the attractive and repulsive effects described by Fritsche and colleges (2020) (Moon and Kwon 2022; Sadil, Cowell, and Huber 2021).

Trying to ascribe biases to past responses is further complicated by *context-independent* biases (e.g., *cardinal bias*) (Fritsche 2016). When sorting trials as a function of the previous response (*resp*_*N-1*_), the sorting variable (*∆R = resp*_*N-1*_ - *stim*_*N*_) is dependent on the physical stimulus feature (*stim*_*N*_) in the presence of cardinal bias. This is in contrast to analyzing stimulus biases where (for an independent stimulus sequence) the sorting variable is independent of the physical stimulus identity *∆s=(stim*_*N-1*_ - *stim*_*N*_*)⊥stim*_*N*_. As a result, any *context-independent* bias, such as repulsion from the cardinal axes, can lead to a dependence of *resp*_*N*_ on *∆R*. This dependance may be why past studies have shown a spurious attraction to future or shuffled trial sequences – an observation that lacks a reasonable causal explanation (Pascucci et al. 2019). Thus, observing a spurious response bias to future or shuffled sequences raises the concern that *any* measured response bias (e.g., even towards the previous trial, *∆R*_*N-1*_) could also be influenced by the same artifact. In Pascucci et al. (2019) and other studies that followed, this issue was addressed by subtracting the average *context-independent* bias from either participant responses or response errors. This method of correction is reasonable, but may actually be insufficient given other *context-independent* anisotropies (e.g., the *oblique effect*) as noted by others (Fritsche 2016). Thus, to reconcile these seemingly paradoxical findings, an analytic framework is needed to successfully disentangle the relative contribution of perceptual, motor, adaptation, and *context-independent* factors.

To address these concerns, we created a model observer exhibiting either stimulus or response driven biases from the previous trial. For parsimony, we will only explore orientation stimuli that feature *cardinal biases* along with the *oblique effect* in this study, but our approach should generalize to other stimulus types (e.g., spatial location, numerosity, pitch). We found that some techniques can reliably distinguish between stimulus and response biases across a range of conditions, but that care needs to be taken to correct for *context-independent* biases. We additionally apply these techniques to an orientation working memory dataset and demonstrate that the history biases observed are primarily attributable to past responses, not to the physical stimulus features. All data and code to implement and expand on these simulations, including power analyses and our analyses of an empirical dataset are available at: https://github.com/TimCSheehan/historyResponseModeling.

## Methods

### Generative Model

To better understand how different sources of bias will ultimately shape behavioral responses, we built a model designed to mimic response properties of human observers. First, we generated an independent and identically distributed (IID) stimulus sequence that uniformly sampled a circular 0-180° feature space (e.g. orientation space). When the sequence is encoded, Von Mises distributed perceptual variability is introduced such that the probability of perceiving a stimulus is governed by the following distribution:

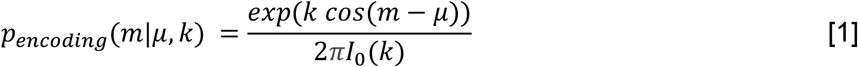

where k and μ are the precision and center of the von Mises distribution respectively, and m is the encoded orientation. I_0_(k) is the Bessel function of the first kind of order 0. We utilize two types of encoding processes. The “biased encoder” features both the *oblique effect*, such that precision is higher around vertical and horizontal stimuli

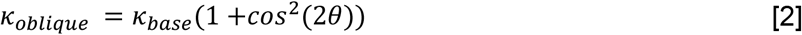

where θ is the stimulus orientation spanning [0, π] and *cardinal bias* such that responses are biased away from the cardinal orientations

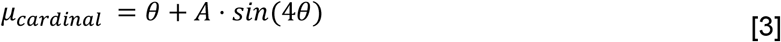

where A=10 is the amplitude of the bias (see Figure 1, *Cardinal Bias* for a depiction of both functions). Note that both *κ*_*oblique*_ and µ_*cardinal*_ have two peaks/cycle as the cosine function is squared for the *oblique effect*. The second encoding model, termed the “uniform encoder”, has constant precision across feature space (κ_uniform_=1.5⋅κ_base_, equalizing average precision) and is centered on the true stimulus value (*µ* = θ).

**Figure 1:**
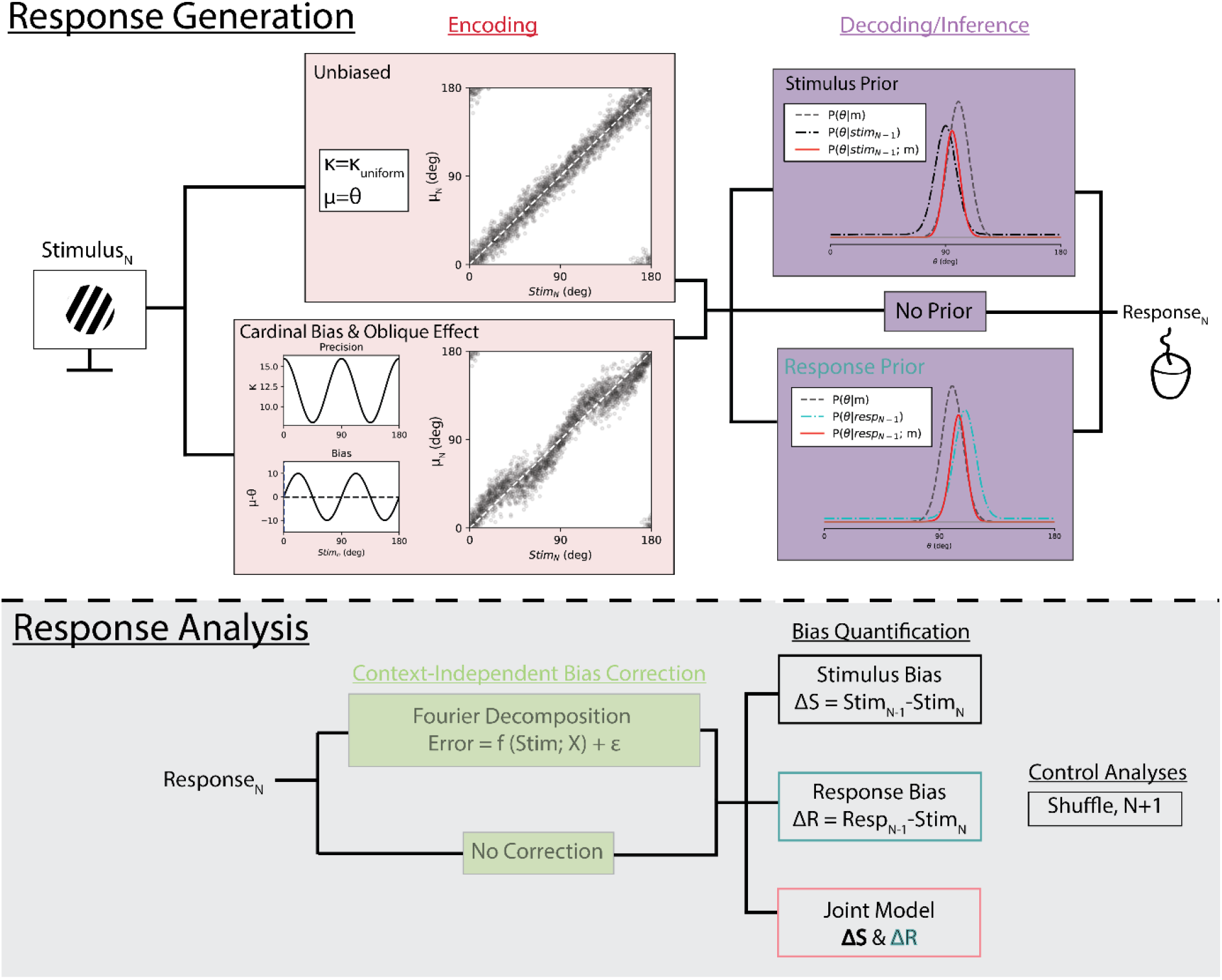
**Response Generation**, on each trial a stimulus is encoded by a biased or unbiased encoder. The encoded representation is interpreted at the inference stage by introducing either a stimulus, response, or no prior for stability. The output from this stage is the response we analyze and used to bias future reports. **Response Analysis**, responses are first corrected (or not) for context-independent biases by fitting a Fourier-like function. We then analyze errors as a function of either the previous stimulus, response, or both. We perform additional control analyses by shuffling trial order or examining the influence of future responses.

On each trial, a random draw from the probability distribution *p*_encoding_ is used to generate a point stimulus estimate m_n_ which is then used as μ in either the biased or the uniform encoding model. This μ parameter, along with the concentration parameter k of the von Mises distribution, generates a probability distribution function (PDF) that defines the stimulus likelihood function^1^. This likelihood is then multiplied by a Bayesian prior centered on either the previous stimulus (“stimulus bias”) or the previous response (“response bias”, Figure 1, *Bayesian Inference*). This prior is based on measurements of natural videos and is a mixture of a von Mises and a uniform distribution to account for both stable random changes across time (van Bergen and Jehee 2019; Felsen et al. 2005). The relative influence of stable and random changes is controlled by the parameter *p*_*stable*_ such that

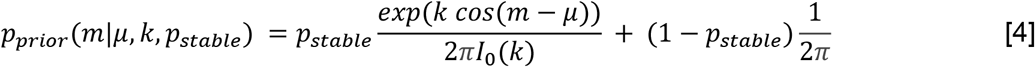

where *µ* is the stimulus or response on the previous trial and κ is constant (building on previous findings suggesting uncertainty on the previous trial does not appear to shape serial dependence in a Bayesian manner (Ceylan, Herzog, and Pascucci 2021; Fritsche 2016; Gallagher and Benton 2022). The maximum value of the resulting posterior

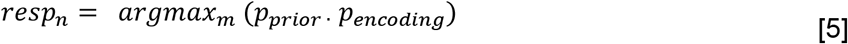

is taken as the Bayes optimal single trial estimate of the stimulus (Figure 1, *Bayesian Inference*, sold line). We equate the output of the model with the “perceived” stimulus value that the participant would indicate with a behavioral response.

### Behavioral Analysis

#### Independent Bias Parameterization

To analyze the results from these different encoding and decoding processes, we sorted response errors as a function of the previous stimulus *(∆S = stim*_*N-1*_ - *stim*_*N*_) or as a function of the previous response *(∆R = resp*_*N-1*_ - *stim*_*N*_). We visualized the resulting bias for each participant by taking a sliding circular mean of the errors as a function of ∆S or ∆R. To simulate typical trial counts of a psychophysics experiment, we ran experiments of 30 participants completing 360 trials each. The magnitudes of history biases were estimated by fitting a derivative of von Mises (DoVM) function:

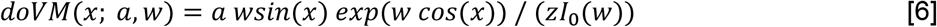

with amplitude, *a*, and width, *w (Sadil et al. 2021)*. These parameters were fit to minimize the RSS errors when *x* corresponds to either *∆S* or *∆R. z* is a normalizing constant such that the amplitude, *a*, corresponds to the height of the resulting function. We additionally performed all analyses using the more commonly utilized derivative of Gaussian function and found similar results, but prefer the DoVM function as it is continuous at ±π.

#### Long-term Bias Correction

Previous studies have attempted to account for any confounds introduced by *context-independent* biases by subtracting out the average bias from either the responses (*resp*_*N*_) or the errors (*resp*_*N*_ - *stim*_*N*_) (Fritsche 2016; Moon and Kwon 2022; Pascucci et al. 2019; Sadil et al. 2021). We perform this correction by first fitting an n=6 parameter Fourier-like decomposition

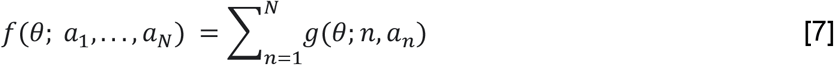

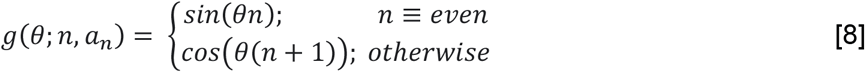

to subjects errors as a function of *stim*_*N*_ and subtracting this function from either the responses (*response correction*: *resp*_*residual*_ = *wrap*(*resp*_*N*_ -*f*(*stim*_*N*_))) or from the resulting errors (*error correction*: *E*_*residual*_ = *wrap*(*resp*_*N*_ -*f*(*stim*_*N*_) -*stim*_*N*_)). Note that correcting responses additionally influences the errors as they are calculated using the modified responses. When analyzing response biases, both corrections impact errors (y-axis) (as correcting responses also corrects errors) while response correction additionally impacts sorting of trials (x-axis). While these two forms of correction ultimately yield similar results, it is important to consider how response correction procedures change the interpretation of any resulting bias (see Discussion).

One concern that arises with analyzing response biases, and a primary motivation for this study, is the presence of ‘spurious serial dependence’ whereby sorting responses as a function of ∆R can give the appearance of attractive biases to the N+1 stimulus or after shuffling the stimulus sequence (Pascucci et al. 2019). As we do not expect the response on a future or random trial to influence our error on the current trial, the presence of such a bias is concerning and may suggest a bias measured relative to past/future stimuli is an artifact of the analysis procedure. To better understand this phenomenon, we additionally consider our errors relative to both the N+1 stimulus and relative to the N-1 stimulus of a shuffled trial sequence.

#### Joint Bias Parameterization

Recent studies have simultaneously modeled the impact of the previous stimulus and previous response (Moon and Kwon 2022; Sadil et al. 2021). We implemented this by parameterizing two DoVM functions modulated by *∆S* and *∆R* and optimized to minimize the residual SSEs. Specifically, we have two vectors *∆S* and *∆R* which are inputs to two DoVM functions. The resulting minimization function is

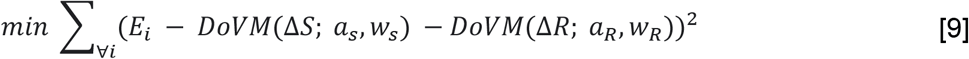

where *E*_*i*_ corresponds to the actual error, *wrap(resp*_*i*_ - *stim*_*i*_), on the *ith* trial.

#### Statistical Analyses

When bias curves are visualized, we include the results of one-sample and paired two-tailed t-tests without correction of the amplitudes of fit DoVM functions.

#### Power Analysis

We performed power analyses to estimate the probability of detecting a significant effect (α<.001) for an experiment conducted with n=30 participants and defined effect sizes and trial counts. For a given experiment, we present the probability of rejecting the null hypothesis that stimulus or response biases are significantly greater and in addition that the magnitudes of the two effects are different from one another.

#### Additional controls

Most experimentalists interested in studying serial dependence intentionally utilize stimulus sequences with a roughly uniform distribution of trial-by-trial stimulus transitions (e.g., *P(∆S)* is uniform). For a variety of factors including inadequate randomization due to low trial counts or the introduction of intentional structure into the distribution, this assumption is often violated to varying degrees (Chopin and Mamassian 2012; He et al. 2010; Maus et al. 2013). To determine how non-uniform stimulus sequences affect measurements of serial dependence, we additionally simulated an analysis pipeline using sequences that feature positive (+) and negative (-) autocorrelations.

The fundamental concern that motivates including simulations with autocorrelated stimulus sequences is that studies attempting to reveal attractive biases to past stimuli or responses may instead only reveal artifacts of their analysis techniques where no biases are present. To assess these concerns, we additionally generate responses where neither stimulus or response serial dependence were implemented to provide a ground-truth case where no biases should be observed (see Figure 1, decoding).

To account for the possibility of a repulsive bias from the stimulus itself, for some experiments we inserted an additive DoVM repulsive bias centered on the previous stimulus with width 1 and variable amplitude.

#### Psychophysical Study

18 participants completed between 192 and 488 (380 ± 15.2, mean ± SEM) trials of a delayed orientation report task. All participants provided informed consent, had normal or corrected to normal vision, and were compensated either in course credit or at a rate of $10/hour. Participants were instructed to fixate on a black fixation cue that was present at the center of the screen 0.5° (degrees of visual angle) and was visible throughout the entire experiment. The trial began with a 1500 ms ITI featuring only the fixation point. Then, two foveally presented oriented gratings subtending 1.5 to 23° degrees of visual angle were presented in succession separated by a 1000 ms inter-stimulus-interval (ISI). Each stimulus had a randomly oriented grating (2 cycles/°, 0.8 Michelson contrast) that was smoothed by a 2D Gaussian kernel with σ=0.5°. Each stimulus was presented for 1s and reversed phase every 125 ms. Each stimulus was followed by a 250 ms filtered noise mask [f_low_=0.25, f_high_=1.0 cycles/°] that changed once after 125 ms. After the second item, a retro cue (the numbers ‘**1**’ or ‘**2**’) indicated the target most likely to be probed (80% validity). On 1/6th of trials a neutral (‘**X**’) was presented in lieu of a retro cue (both items equally likely to be probed). The retro cue was followed by a blank delay period 2500 ms. Participants then controlled a black response dial (using the “ASDF” buttons on a standard QWERTY keyboard) and they were given between 500 and 5000 ms to match the orientation of the probed stimulus. After pressing the space bar to confirm their response or timing out, the dial disappeared, and feedback was provided for 2000 ms by displaying the unsigned error in degrees and turning the response dial green if participants were closer than 10° and red otherwise.

## Results

### Serial Dependence Without Cardinal Bias

We first analyzed responses in a model without *context-independent* biases featuring either stimulus serial dependence, response serial dependence, or no serial dependence (columns *Left, Center, Right* respectively, Figure 2). For this simulation, and unless otherwise noted, we use κ_base_ = 8 and therefore κ_uniform_ = 12. The first row shows biases relative to the previous stimulus and reveals that trials with true stimulus bias (Figure 2A) show a larger stimulus (*∆S*, black curve) relative to response (*∆R*, teal curve) bias. We additionally observed a larger response bias when the underlying source of the bias is towards the previous response (Figure 2B). Together, this suggests that, in the absence of *context-independent* biases, the relative magnitudes of stimulus/response serial dependence is a good proxy for the dominant source of the bias.

**Figure 2:**
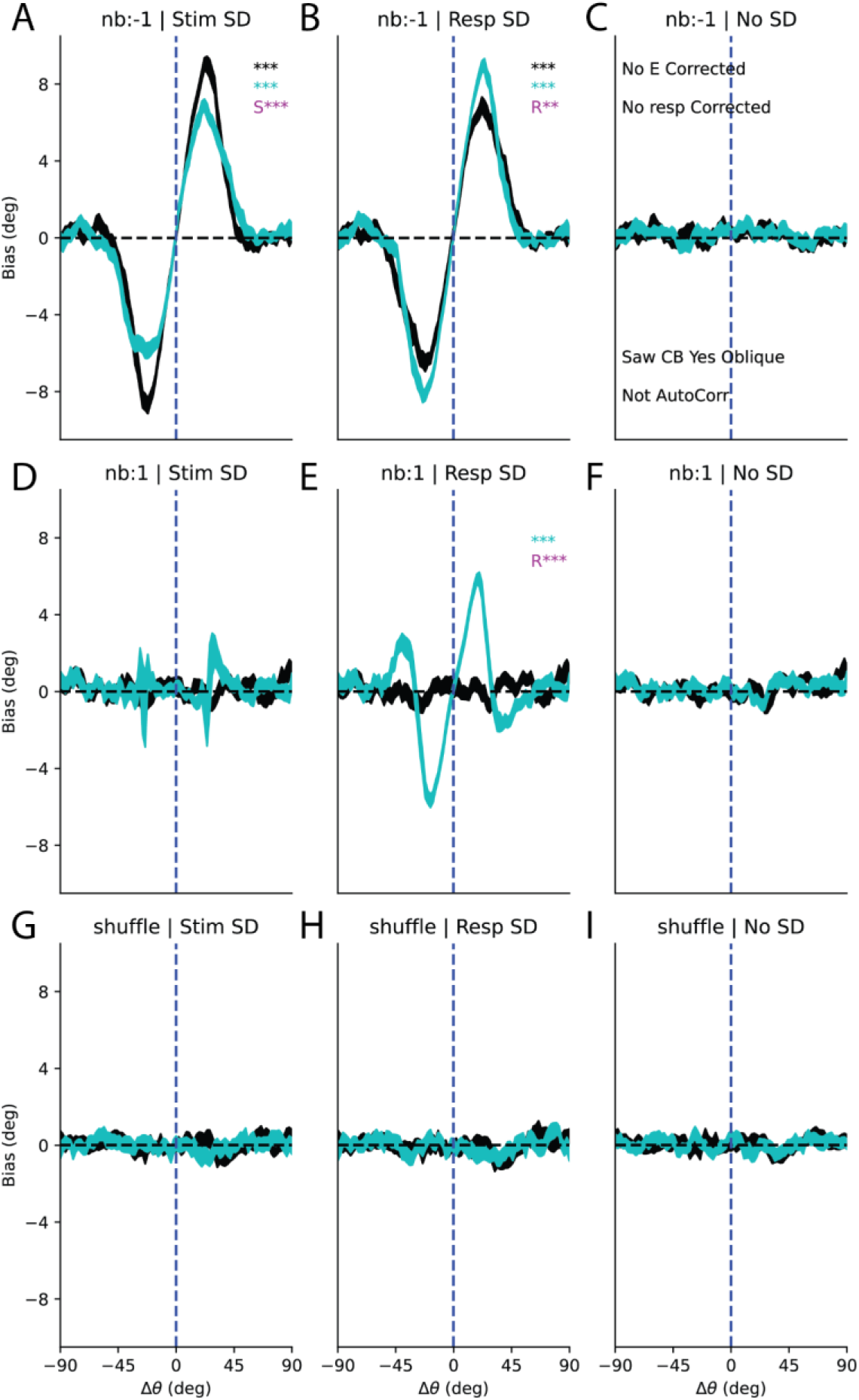
Stimulus (black) and response (teal) bias curves for all response simulations. (Left, A,D,C) column corresponds to responses generated with an attraction towards past stimuli, (center, B,E,H) column features responses attracted towards past responses, and (right, **C**,**F**,**I**) column has no history biases. (Top, **A-C**) row computes ∆θ relative to previous trial, (middle, **D-F**) row computes ∆θ relative to future trial, and (bottom, **G-I**) row computes ∆θ relative to the previous trial after shuffling the stimulus order. Both **A** and **B** show significant attractive biases towards past stimuli and responses with larger attractive biases towards the underlying source of the bias. We additionally observe an attractive bias towards the future response **E** that is an artifact of our sorting procedure. *, p<.05; **; p<.01; ***, p<.01, Bonferroni corrected for 9 stimulus conditions; R, response bias significantly greater than stimulus bias, S, stimulus bias significantly greater than response bias.

Critically, the only artifactual bias occurs when examining *∆R*_N+1_ when there was a genuine bias response bias (Figure 2E). This demonstrates that cardinal or other history independent biases are not necessary to observe artifacts in analyzing response biases in the presence of true response dependence and suggests that such an artifact is an indicator of a bona fide bias in the data. We explore why this N+1 artifact arises in the next section.

### The N+1 response bias artifact

Ensuring that there is no bias toward future responses (i.e. the *N+1 trial*) has been suggested as a valuable control when evaluating response biases (Pascucci et al. 2019). However, as noted above, we find an attractive bias when sorting trials by *∆R*_*N+1*_ when there is a true response-based serial dependence effect. To understand why this bias occurs, we first identified an important distinction between sorting trials based on the past versus future response. Importantly, *resp*_*N-1*_ is independent of *stim*_*N*_ and accordingly *P(∆R*_*N-1*_) is uniform (Figure 3A). However, *resp*_*N+1*_ is *not* independent of *stim*_*N*_ because it is influenced by a prior centered on either *stim*_*N*_ or *resp*_*N*_ (depending on the source of the bias) resulting in a highly non-uniform distribution (Figure 3A, *P(∆R*_*N+1*_)). To explore why the *∆R*_*N+1*_ spurious bias occurs, we considered two possible outcomes on the current trial, an error *CW* or *CCW* relative to the true stimulus. For the purposes of this visualization, we used the average absolute error of our unbiased observer, 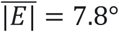. For observers exhibiting response-based history biases, these *CW/CCW* errors generate distinct priors (Figure 3B) that differentially shape future responses. These priors shift *P(∆R*_*N+1*_) towards the current response (Figure 3C). The difference in relative probabilities of the previous response error multiplied by the average response error 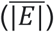 perfectly captures the measured “spurious” response bias (Figure 3D, 2E). Thus, spurious biases measured by examining the influence of the *N+1* response are *expected* if the underlying source of the bias is a prior centered on the preceding response. Because of this, examining the *N+1* influence is not a pragmatic control analysis and researchers should instead opt for a shuffled trial sequence which does not exhibit spurious biases when response biases are present in a dataset.

**Figure 3:**
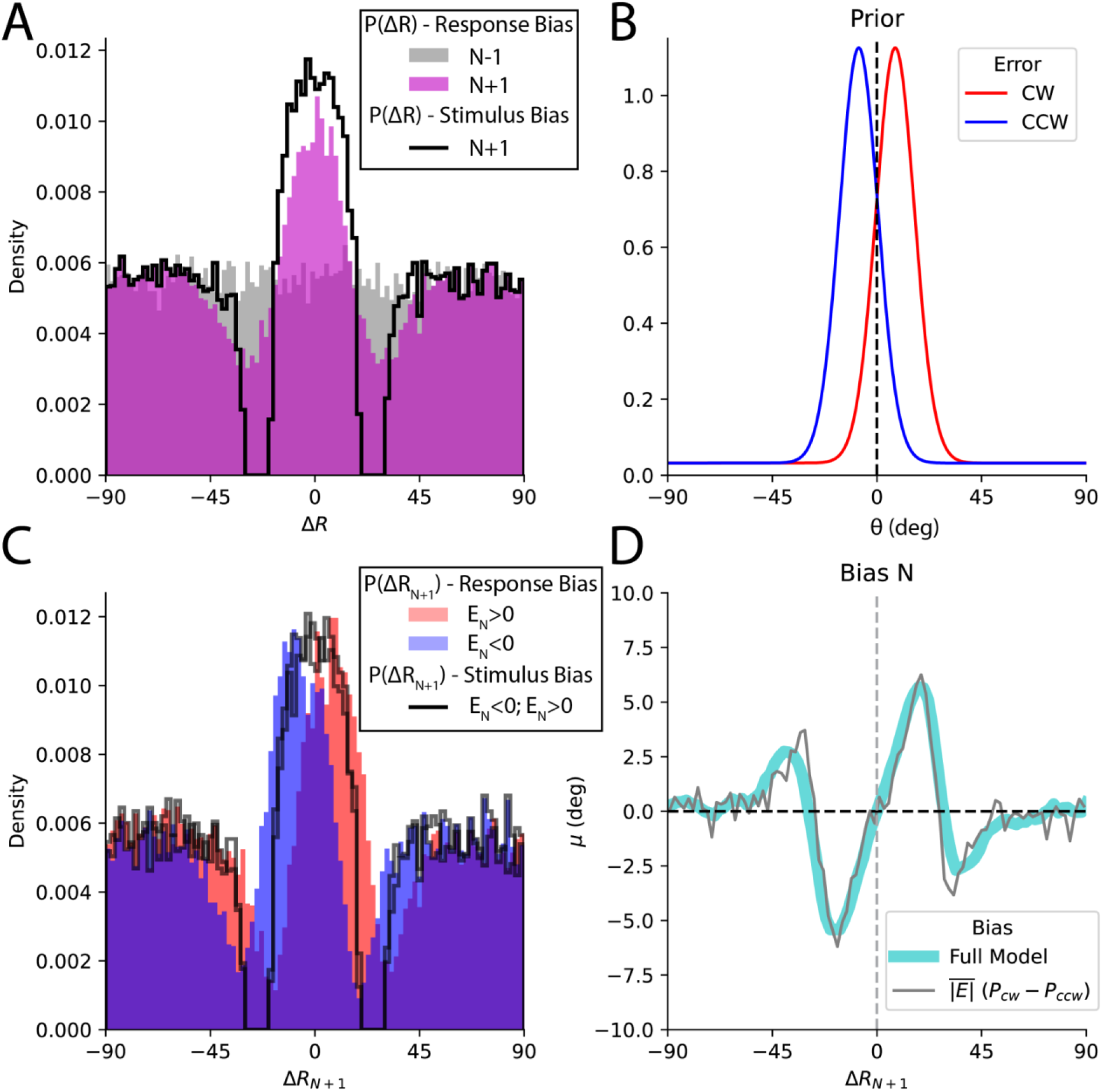
**A**. P(R_N-1_), gray, is uniform but P(∆R_N+1_), magenta, shows an overrepresentation for small changes. Additionally shown is P(∆R_N+1_) for stimulus serial dependance (black trace). **B**. hypothetical priors following a misperception of the average magnitude for our model (7.8°) in the CW or CCW direction. **C**. P(∆R_N+1_) on trials with CW or CCW misperceptions are shifted relative to each other. This shifting does not occur when the bias source is the stimulus instead of response (black traces) **D**. The average (unsigned) error multiplied by the difference in the P(∆R_N+1_) for CW and CCW responses captures the measured spurious bias.

### Serial Dependence with Context Independent Biases

We next analyzed serial dependence after additionally including cardinal bias and the oblique effect at encoding. Both the precision κ and expected value μ were modulated by the stimulus identity resulting in an encoding process that showed characteristic bias and variance patterns of *cardinal bias* and the *oblique effect* (Figure 1). The result of this biased encoding process was then modulated by the same Bayesian prior as used in the previous section. When analyzed, the resulting responses show an increased response bias and a substantial ‘spurious’ response bias in the absence of any history biases (Figure 4A-C) demonstrating that *context-independent* cardinal biases can introduce an artifact as suggested previously (Fritsche 2016; Pascucci et al. 2019).

**Figure 4:**
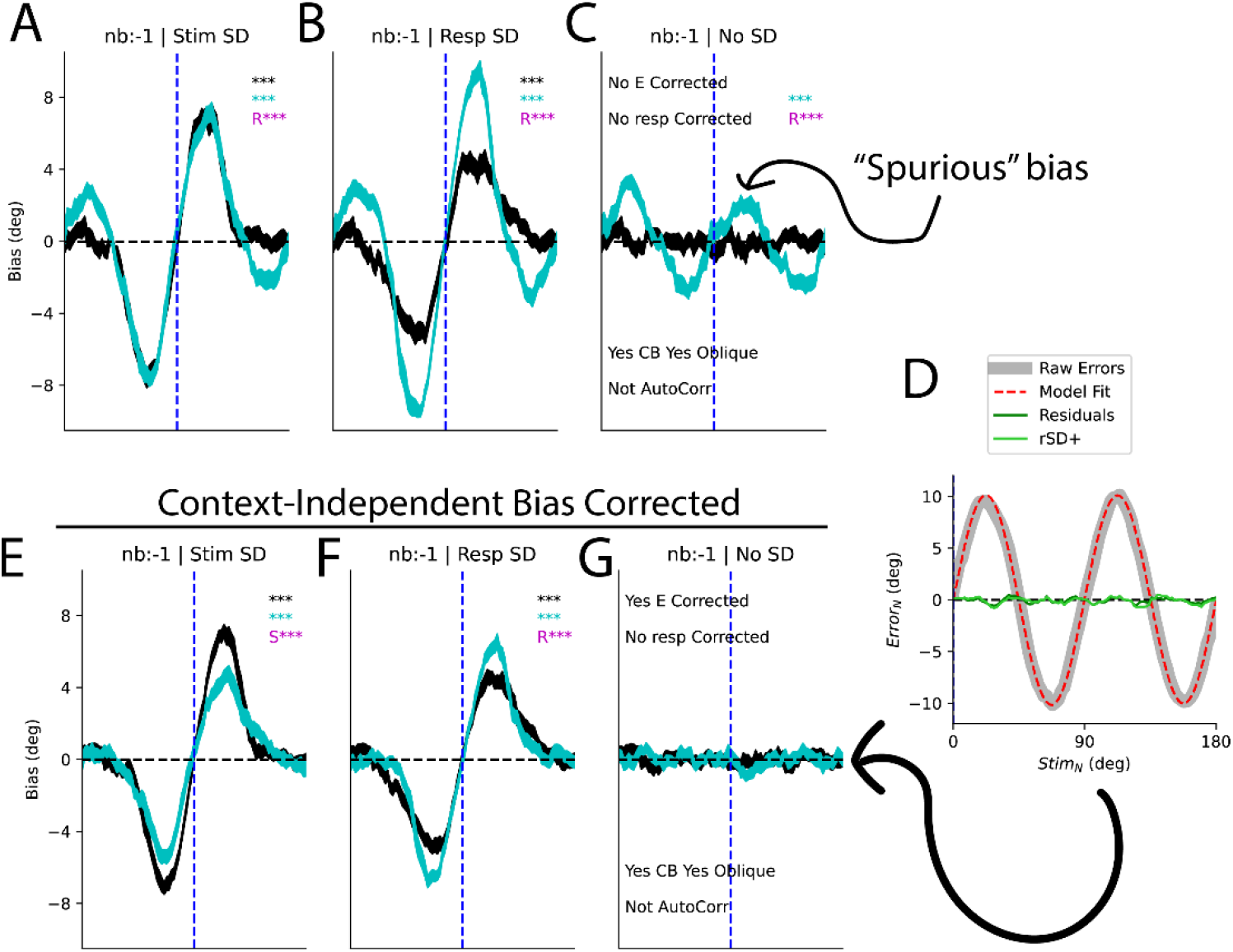
**A-C**. Response/stimulus biases computed using the raw errors results in a spurious response bias (see Fig S1 for all bias curves) **D**. Context-independent biases can be corrected for by fitting a model to responses such that the resulting residuals are not biased as a function of stimulus identity. The light green trace (rSD+) is the residuals when history dependent bias (serial dependance) is present when fitting the history independent bias model. **E**-**G** Response/stimulus biases computed using the residualized errors.

This confound is more concerning than the *∆R*_*N+1*_ bias we found in the previous section because an attractive response bias is found even when no underlying serial dependence is present in the generated data (Figure 4C) or when trial order is shuffled (Figure S1A). Previous studies have tried to address this bias by regressing out the stimulus specific bias from either the errors or the responses. This has generally been achieved by fitting either a higher order polynomial or sinusoidal function to the raw data. For the purposes of this study, we utilized a 6-parameter Fourier like composition of sine/cosine functions of varying frequencies which is more flexible (see eq. 7). Our use of circular functions avoids edge effects found with polynomial fits. We fit this function to errors and subtracted the best-fit function to correct for these biases (Figure 4D, *red dotted-line*). This correction substantially reduces any trace of systematic biases (Figure 4D, green). We opt to correct errors, but not responses, as this allows *∆R*_*N-1*_ to reflect the relative location of the previous response.

Correcting for *context-independent* biases in response errors appears to completely remove the presence of spurious biases and returns the relative magnitudes of biases to what is expected given their respective sources (Figure 4E-G, See Figure S1 for bias curves corresponding to shuffled and N+1 trials). This is critical as this regression-based approach is an effective way to correct for *context-independent* biases and ensure the presence of measured response history biases is not just an artifact. This correction process does nothing to account for differences in variability as a function of the stimulus (the *oblique effect*) but still removes any trace of artifactual responses in the shuffled condition. We separately analyzed the influence of autocorrelations in the sequence of stimuli presented and found no evidence that they introduce new artifacts (Figure S2).

### Cardinal biases cause spurious response biases

It is not surprising that introducing biased stimulus representations could introduce cofounds. In a general sense, this is because *Error*_*N*_ is dependent on *stim*_*N*_ and furthermore *∆R* is no longer independent of the absolute stimulus value. Why this leads to spurious history biases is not particularly intuitive, so we provide a brief demonstration here. First we visualize the joint distribution *P(Stim*_*N*_, *∆R)* which shows the two variables are clearly not independent (Figure 5A). Note that we are not specifying which trial is the inducer (eg. *N-1/ N+1*) as this spurious bias is unchanged even after shuffling trial order. The conditional distributions *P(Stim*_*N*_ | *∆R)* for two subsets of ∆R reveal how dramatically *P(Stim*_*N*_) is interdependent on *∆R* (Figure 5B). We can then approximate the predicted spurious bias as the dot product of the normalized rows of *P(Stim*_*N*_, *∆R)* with *μ*_*cardinal*_ *(Stim*_*N*_) (Figure 5C, 4A) to get the expected bias

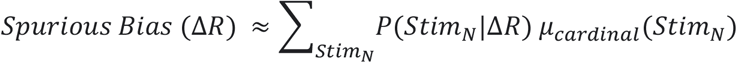

(Figure 5D, black). This process captures the “spurious” response bias from the shuffled response distribution (Figure 5D, teal). Note that when sorting trials based on the previous stimulus instead of responses, *P(Stim*_*N*_|*∆S)*, is independent and does not give rise to spurious history bias.

**Figure 5:**
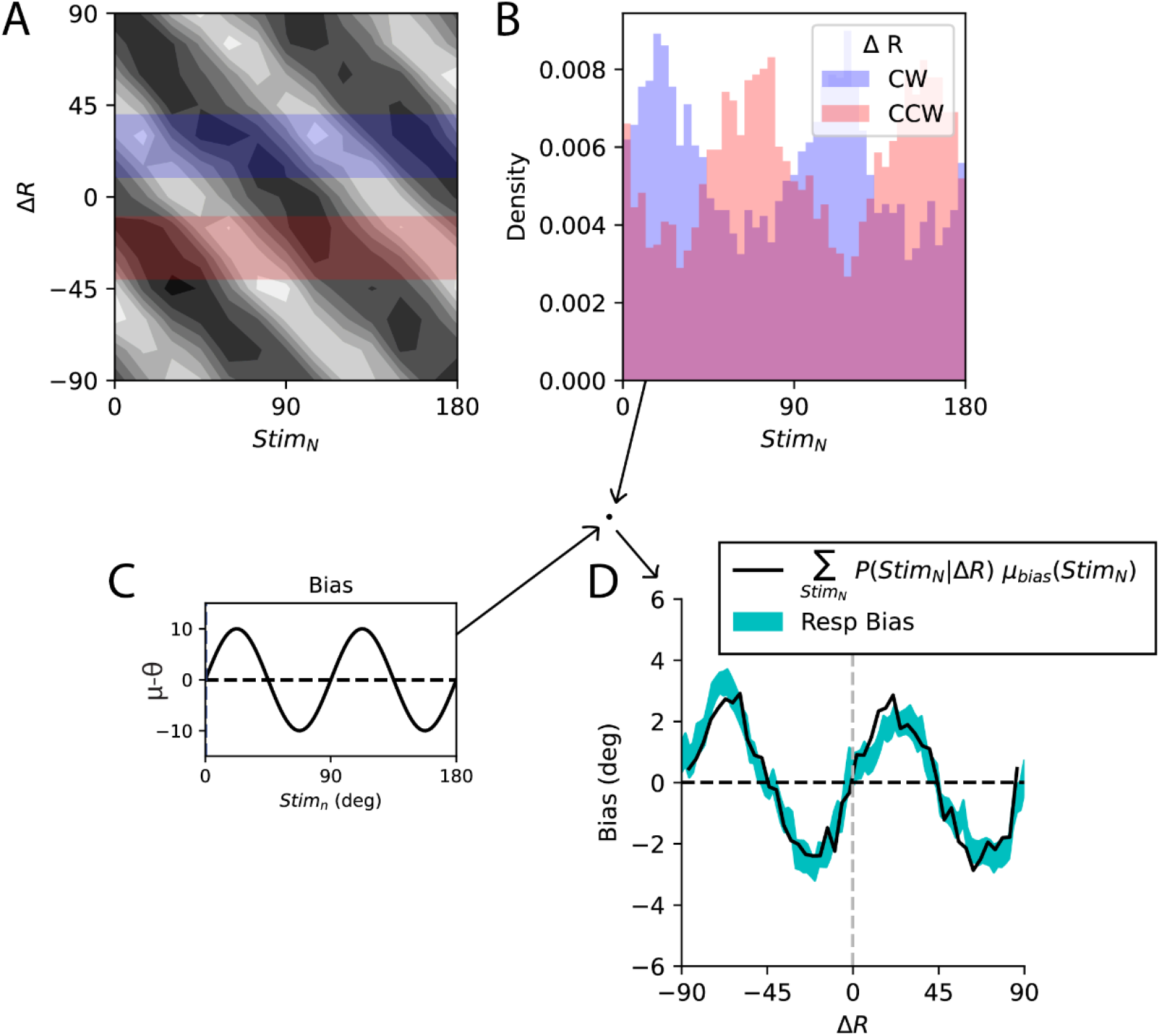
**A**. The distribution of ∆R is not independent of Stim_N_. **B**. We illustrate the distribution of Stim_N_ for the subsets of trials highlighted in (A). **C**. Expected error as a function of Stim_N_. **D**. Response bias (teal± SEM) is captured by the product of P(Stim_N_|∆R) and μ(Stim) (black).

### Simultaneous modeling of stimulus and response

Two recent studies have tried to disentangle the relative contributions of stimulus and response history biases (Moon and Kwon 2022; Sadil et al. 2021). Using this approach, the two functions are fit simultaneously instead of fitting a single two parameter function separately to *∆S* and to *∆R*. Theoretically, this should better disentangle the sources of the bias and the approach has revealed the surprising possibility that stimuli could simultaneously be *repelled* from the previous stimulus but have an even larger attraction to the previous response (Moon and Kwon 2022; Sadil et al. 2021). This approach is interesting but may be problematic as the two regressors are highly collinear, which poses a challenge for interpreting the fit parameters. We applied this approach to two simulated datasets, our full model featuring cardinal bias and correction for that bias, and a new model which introduces repulsion from the previous stimulus (see Methods, *Joint Bias Parameterization)*. First, we visualized the average individual fits to our corrected errors (as presented in Figure 4C) and note that while our modeling approach correctly captures the predominant bias source, the non-causal source is still of a similar magnitude (Figure 6A, left). When we apply our joint fitting procedure to the same data, we are better able to capture the true underlying source of the bias (Figure 6A, right). To compare the effectiveness of these alternative approaches, we conducted a power analysis for detecting significant biases while varying trial counts and precision (see Methods). First, we note that our power to distinguish between stimulus and response biases was higher for low precision participants across model types (Figure 6B). Critically, however, we note that the independent model consistently detects a significant effect of the non-inducing feature (Figure 6B, *top*) while the joint model is much less likely to detect a significant non-causal effect (e.g., Figure 6B, *bottom, ∆S* is close to 0% power for the joint model given true response serial dependance). This suggests the joint model is better powered to avoid Type II errors. See Figure S4 for a power analysis further broken down by trial count.

**Figure 6:**
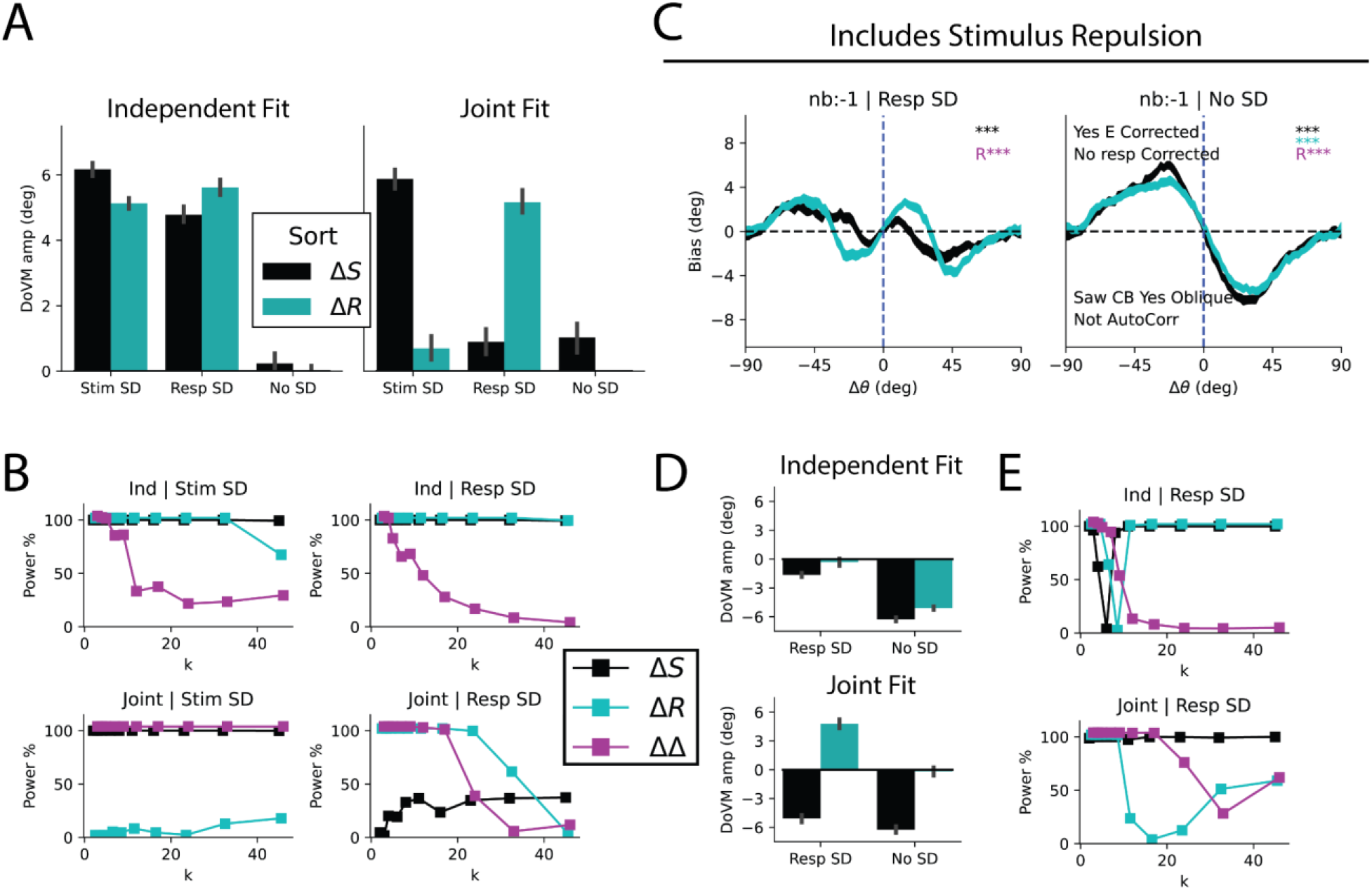
**A**. Fit magnitudes for independent and joint model fits. **B**. Power analysis across a range of k values for independent and joint models. Power is the % chance at finding a significant effect with n=30 participants at α=.001. ∆∆ refers to direct comparison of magnitude of ∆S and ∆R (paired t-test). **C**. Bias curves for an observer featuring stimulus repulsion, additional curves Figure S3. **D**. Joint fit is able to capture magnitudes and signs of true biases while independent model fails to separate the two. **E**. Power analysis reveals challenges in calculating bias magnitudes when the two competing forces are of approximately equal (0 power for ∆R at k=8 for independent model. Expanded power analysis presented in Figure S4.

We next applied the same approach to an observer featuring repulsion from the previous stimulus implemented at encoding to determine how well the joint/independent models captured these opposing effects. This is challenging because stimulus repulsion acts to counteract the influence of response attraction (Figure 6C, Figure S3). We found the joint model was better able to capture the underlying bias source (Figure 6D) and generally had much better power at distinguishing between their influences across a range of conditions (Figure 6E, bottom, Figure S4). This power analysis revealed an interesting phenomenon that may be common in the serial dependance field. For the independent model, particular values of k led to stimulus and response biases that largely counteracted one another leading to 0% power (Figure 6E, top). Importantly, the joint model was able to reliably detect response biases over this same range (Figure 6E, bottom). This idea of opposing attractive and repulsive biases could suggest why null or weak results are common in studies of serial dependance and may provide a new avenue to analyze existing datasets.

### Application to Empirical Data

We conclude by applying the techniques and principles developed above to an existing unpublished dataset. Participants (N=18, 6840 trials total) viewed a sequence of two oriented gratings presented foveally in succession and reported one of the stimuli by rotating a response dial with the keyboard after a 3.5s delay period. This experiment included partially valid retrocues, the full details of which are described in the Methods and schematized (Figure 7A, Figure S5A). We first noted that responses showed strong *context-independent* biases that were non-sinusoidal (Figure 7B, gray). We first attempted to fit *context-independent* biases using a 6 parameter Fourier-like function as with our simulation, but found it was a poor match with large residuals (Figure 7B, *light green*). To fully capture the structure, we instead opted for a 12-parameter version which achieved a much tighter fit and smaller residuals (Figure 7B, *dark green*). We then examined history biases non-parametrically for the N-1 trial with and without shuffling trial order. For the shuffled responses, the correction procedure removes a spurious response bias seen in the raw responses (Figure 7C, bottom). The *In Order* trials show strong stimulus- and response-based biases (Figure 7C, top). We next examined stimulus and response biases both separately and using a joint model. To improve our power, we bootstrapped responses by randomly resampling 360 trials with replacement for 1024 surrogate participants. Participants showed strong attractive biases when sorting by *∆S*_*N-1*_ & *∆R*_*N-1*_ (Figure 7D, left). Critically our correction procedure removed the *context-independent* bias artifact (Figure 7C, bottom-right). Consistent with our previous simulations, we found that response biases were inflated for all analyses and are significantly greater than 0 after shuffling when we didn’t correct for *context-independent* biases (Figure S5C). When quantifying history biases independently, both stimulus and response biases were highly significant, but response biases were significantly stronger (Figure 7D, left, *In Order*). Importantly, we did not observe any stimulus or response biases for the shuffled trial sequence (Figure 7D, left, *Shuffle*). When we applied the joint fitting procedure, we found that only response bias was significantly greater than 0 suggesting that response biases are the dominant source of attractive biases in this data set. We thus demonstrate that our analysis procedure can be applied to empirical datasets and that simultaneously modeling biases can lead to insights otherwise hidden by traditional approaches.

**Figure 7:**
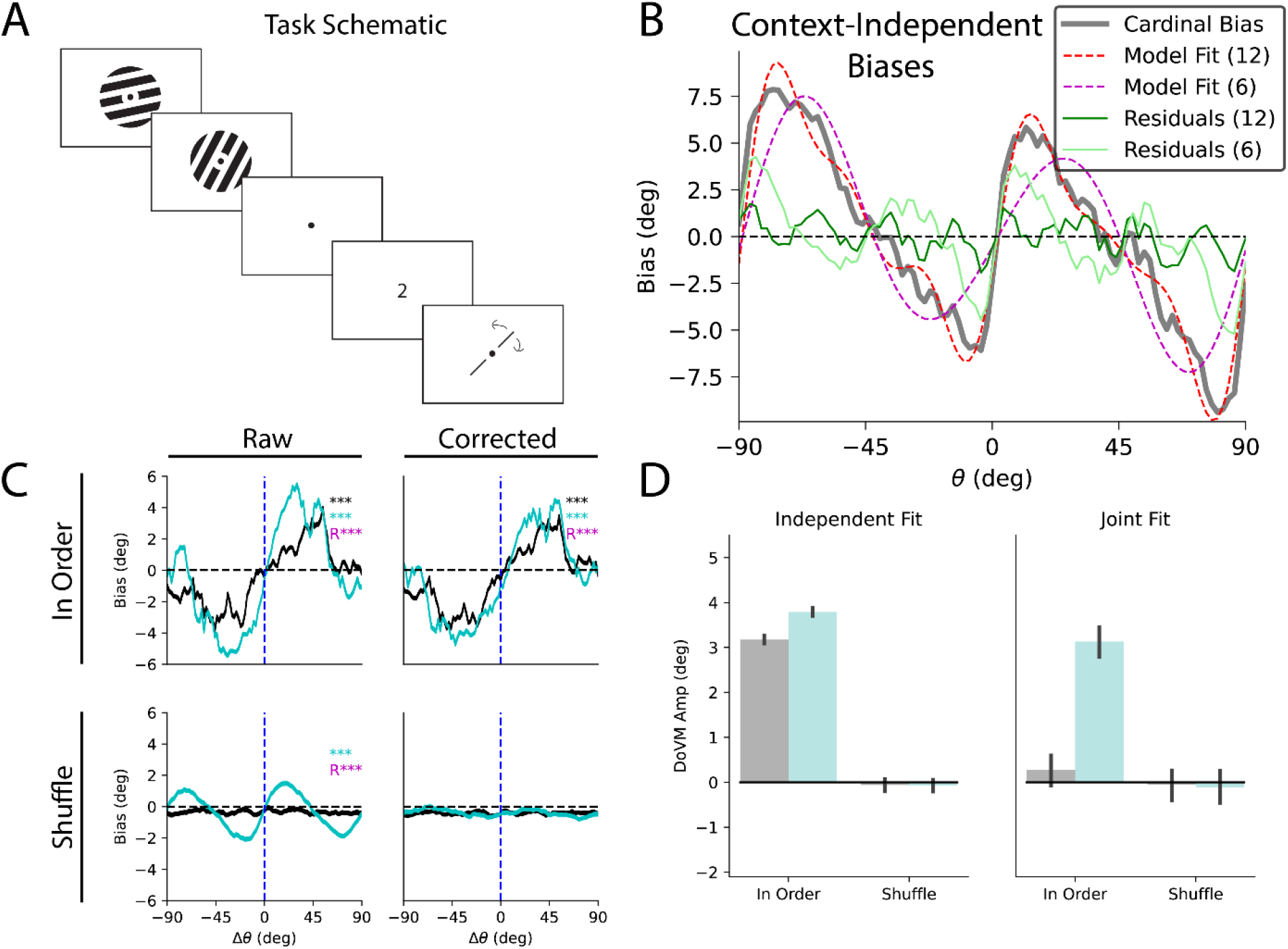
**A**: simplified task schematic. Participants reported 1 of 2 foveally presented stimuli after a delay. **B**: Responses showed strong context-independent biases (gray). These were corrected by fitting a 12-parameter Fourier based parameterization to the pooled errors (red) resulting in unbiased residuals (green). **C**: Top, N-1, both raw and corrected responses show larger biases when sorting by past responses than stimuli; bottom, shuffle, uncorrected responses show a spurious response bias after shuffling trial order (left) that is eliminated after context-independent correction (right). **D**: While the independent model suggests both stimulus and response biases, joint model reveals bias is driven by responses.

## Discussion

The goal of this modeling work was to provide a comprehensive exploration of methods to dissociate stimulus and response biases in the presence of potentially confounding *context-independent* biases such as *cardinal bias*. This work was motivated by an acute interest in analyzing response biases combined with a concern that any bias measured could be an artifact of the analysis procedure. We first recap the lessons from our simulations and then discuss considerations that need to be made when analyzing such biases in empirical studies. Last, we briefly consider the psychological implications of our own empirical findings and recent related work.

We first identified a spurious future bias that is found specifically when sorting by *∆R*_*N+1*_ (Figures 2-3). This bias is only observed in the presence of true response biases and is found in the absence of (or after correcting for) *context-independent* biases. This phenomenon is a signature of response biases and may be interpreted as evidence for previous responses rather than previous stimuli inducing a history bias (and notably this bias does not emerge under stimulus induced biases, Figure 2D). Importantly, there is no analogous spurious future bias after shuffling the trial order before assessing serial dependence (Figure 2H). Thus, the analysis of ∆R_N+1_ biases should primarily be used as a confirmatory step for the presence of response biases rather than a control for the influence of *context-independent* biases.

More problematic are artifacts introduced by *context-independent* biases (e.g., *cardinal bias)*. These can lead to a spurious attraction between shuffled responses (Figure 4C). In our simulations, the spurious response biases were eliminated after regressing out this bias (Figure 4D, G). These biases emerge due to the influence of *context-independent* biases on all responses which is why shuffling does not remove them (Figure 5). When applying this correction procedure to our empirical dataset, the cardinal biases we observed were much steeper than the sine wave used in our simulation and necessitated additional higher frequency components to achieve truly unbiased residuals (Figure 7B). We increased the expressivity of our correction procedure until the errors sorted by *Stim*_*N*_ and *∆R*_*Shuffle*_ were flat and unbiased (ultimately using a model with 12 free parameters). We were then confident that any response biases were genuine and not an artifact. Here, we observed a response bias ∆R_N-1_ that was significantly larger than our stimulus bias ∆S_N-1_ (Figure 7D, *Independent Fit*).

Lastly, we found promising results utilizing a joint modeling approach that was introduced in a pair of recent studies (Moon and Kwon 2022; Sadil et al. 2021). Our analysis of simulated data showed that despite stimuli and responses being highly correlated, the joint approach was generally able to capture the true source of the bias (Figure 6 A, D). The reliability of this approach was greatly improved when participants were less precise and when there were greater trial counts per participant (Figure 6B, E, S4). Applying this approach to our empirical dataset revealed strong evidence for a history bias that originated from responses, not stimuli (Figure 7D, *Joint Fit*). Surprisingly, this response bias continued back many trials offering a new potential interpretation of past studies that have similarly long-acting biases (Figure S5) (Fritsche et al. 2020; Gekas, McDermott, and Mamassian 2019). Our interpretation of this being a response driven bias is strengthened by the fact that other metrics, including the independent fits and the *∆R*_*N+1*_ bias, all aligned closely with metrics observed for our response-driven simulated observer. Thus, simulated observers offer a valuable tool to infer the origin of biases given the outputs of the various metrics we have tested.

Throughout this manuscript, we present stimulus and response driven biases as if they are mutually exclusive. In reality, it is equally, if not more likely, that the inducing feature from the past is the *perceived* stimulus (rather than the response per se). This is supported by past work that has attempted to directly disambiguate perceived from reported orientations (Cicchini et al. 2017) or work that has utilized change detection rather than continuous report paradigms (Fischer and Whitney 2014; Fritsche et al. 2017; Sheehan and Serences 2022). That said, others have shown that attraction is not generated unless a stimulus is reported and that attraction may instead be towards the reported rather than perceived location (Pascucci et al. 2019; Sheehan, Carfano, and Serences 2022). In any case, with continuous report paradigms we often don’t have any means of directly accessing the identity of the perceived stimulus and so we opt here to use the more general term of “response” throughout this paper as the behavioral response is typically the best/only proxy for the internal perceptual representation. Further disambiguating the physical act of responding (and the associated motor/decisional circuits) from the perception of the stimulus will require careful experimental designs or neural measures that can assess internal representations at different stages of information processing. Thus, finding a bias driven by past responses (rather than physical stimulus identity) as we did primarily suggests that attraction is toward a post-retinal representation or transformation of the stimulus. In retrospect this claim may seem obvious, as the brain has no access to the stimulus per se and will always be relying on internal representations that deviate from the original stimulus feature (Eggermont 2007; György Buzsáki 2019; Lettvin et al. 1959).

Now that there are several studies showing strong evidence for response over stimulus driven effects (Moon and Kwon 2022; Sadil et al. 2021), the goalposts have shifted to further disambiguate exactly which response related components are driving these effects. Change detection paradigms or generally un-correlating responses from perception offer promising avenues to explore this possibility further (Braun, Urai, and Donner 2018; Sheehan et al. 2022; Zhang and Luo 2022). That said, we argue here that examining biases just as a function of the physical identity of the previous stimulus is ignoring the important role of other biases in shaping the perception of current and past stimuli and may lead to an under and mismatched measurement of the true underlying bias (Pascucci et al. 2019; Sadil et al. 2021).

In the behavioral experiment we report here, there was no direct correlation between the final response and motor action as the probe was initialized in a random location and was controlled by button presses. Thus, we can likely rule out a purely motor origin for the attractive biases that we observed. The nidus of the attractive effect could instead be residual traces tied to memory maintenance, a distinct circuit directly tied to representing sensory history, or plausibly a sensory effect tied to the response or feedback signal presented at the end of the trial (Akrami et al. 2018; Barbosa et al. 2020). Only through additional experiments and analyses that control for these additional possible sources of perceptual biases can we further refine our understanding of these processes.

By demonstrating that the influence of *context-independent* biases can be reliably corrected for – while simultaneously highlighting the concerns raised if they are not – we hope to guide future endeavors to identify the true source of history biases. In our own experiment, we found strong evidence for an attractive bias centered on the previous response rather than the physical identity of the stimulus. We further found evidence for this attraction extending back 6 trials and separate evidence for a repulsion from the physical identity of the stimulus for trials 2, 3, 5 and 6 trials back. This pattern matches prior observations and supports the idea that the stimulus presentation leads to a repulsive bias at encoding while more high-level decisional representations impose a prior of stability (Braun et al. 2018; Moon and Kwon 2022; Papadimitriou, White, and Snyder 2016; Pascucci et al. 2019; Pegors et al. 2015; Sadil et al. 2021; Sheehan and Serences 2022; Zhang and Alais 2020; Zhang and Luo 2022). Such a framework additionally fits with general frameworks like efficient encoding and Bayesian inference seen in perception (Wei and Stocker 2015) and pattern separation and completion seen in various networks across the brain (Cayco-Gajic and Silver 2019).

## Supplemental Materials

**Figure S1:**
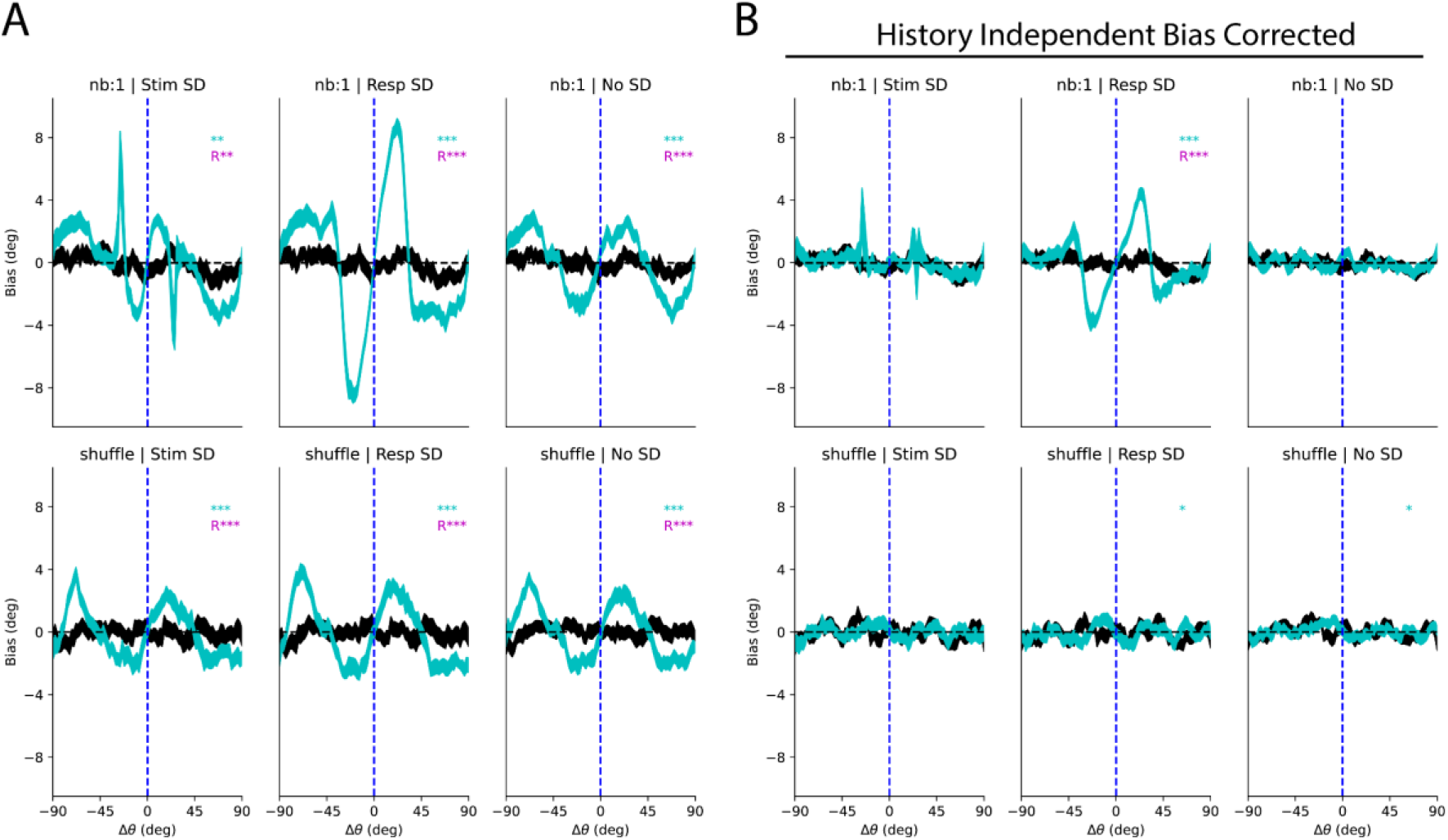
Bias curves for N+1 and shuffled distribution for corrected (**A**) and uncorrected (**B**) errors from Figure 4.

**Figure S2:**
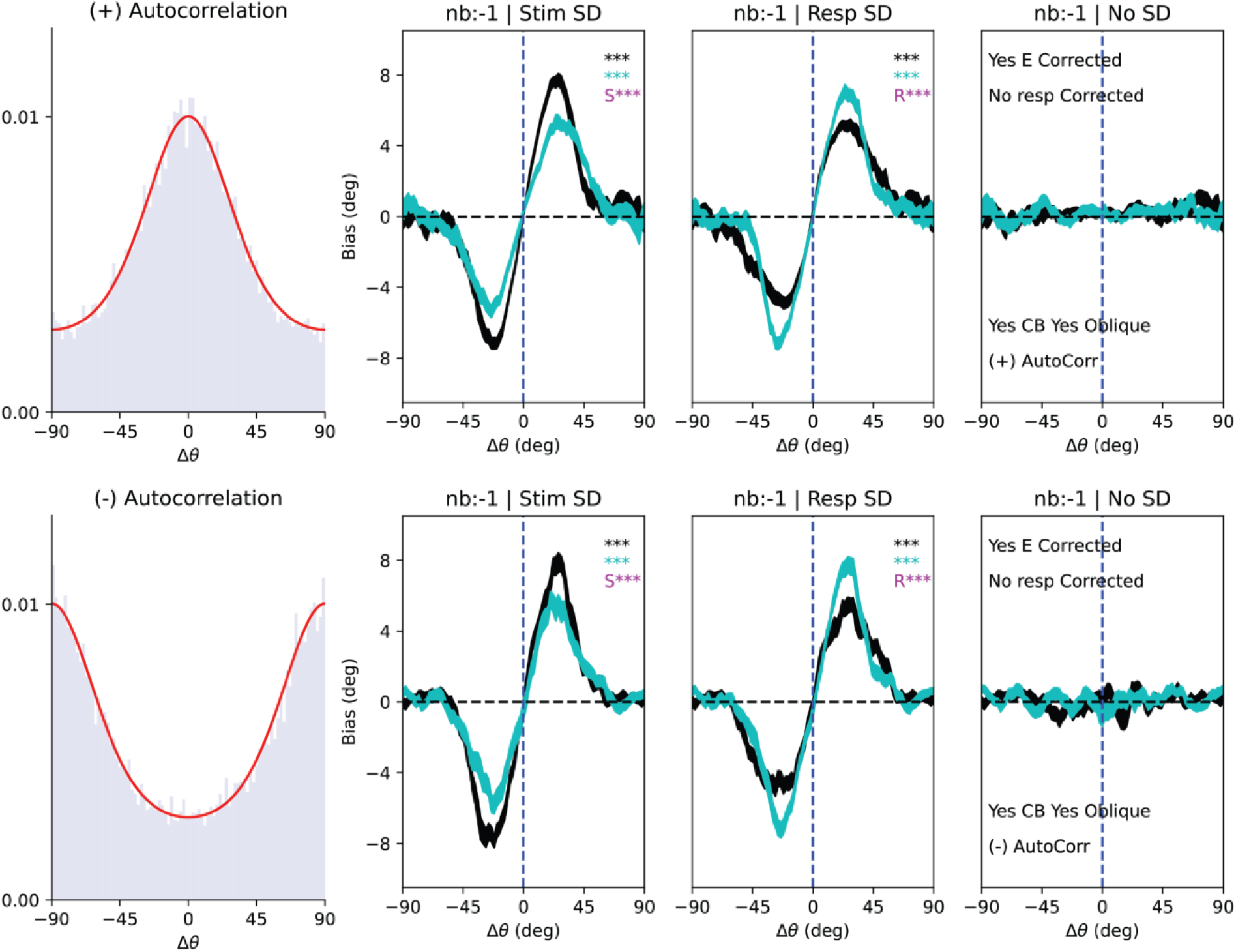
Non-independent Stimulus Sequences. We simulated the analysis of observers where stimulus sequences were non-independent and exhibited strong positive (top left) or negative (bottom left) autocorrelations. Despite the presence of these strong stimulus autocorrelations, their presence alone does not introduce any additional artifacts into our analysis procedure.

**Figure S3:**
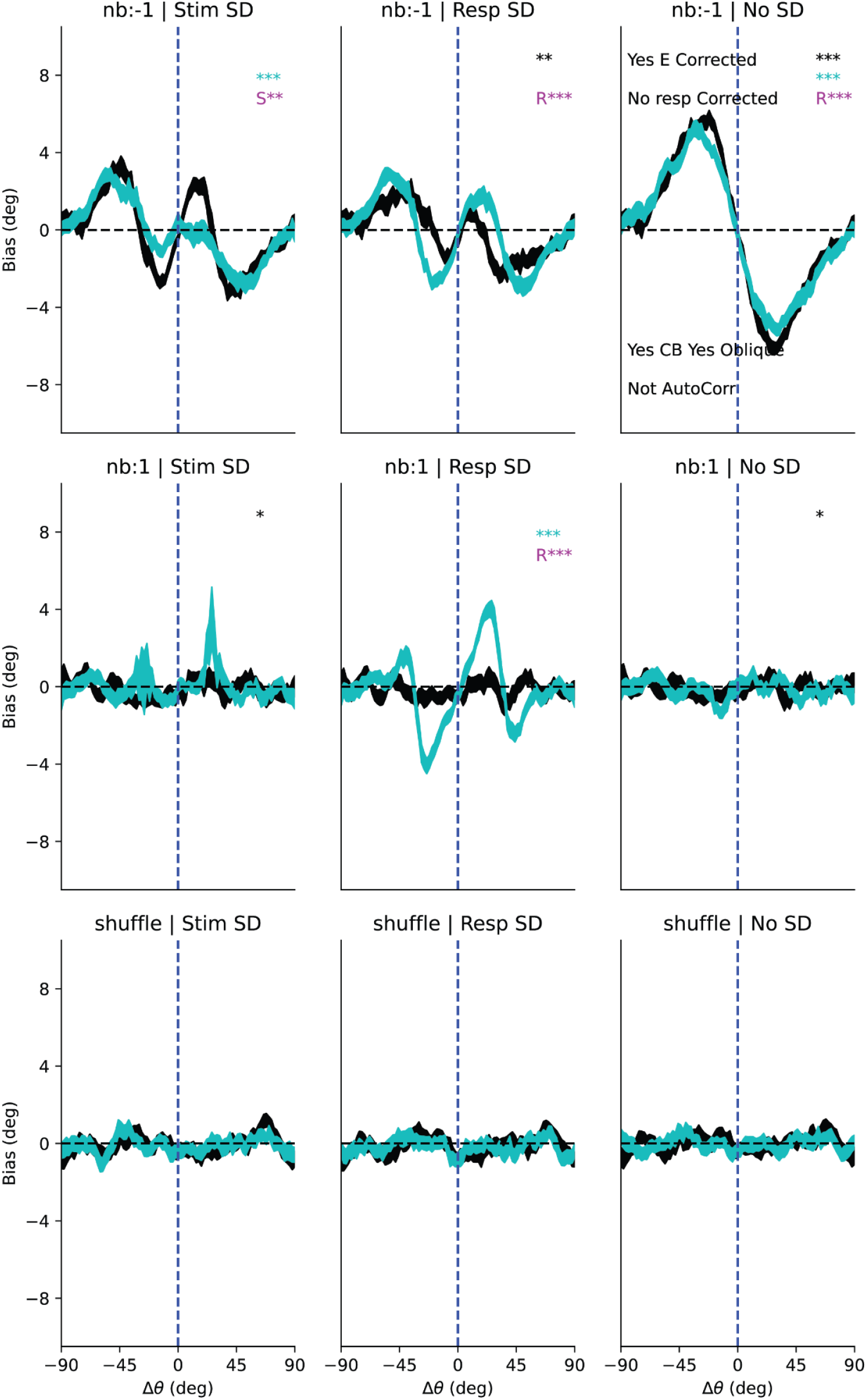
All bias curves for observer with stimulus specific repulsion. Note that the left column is an observer that is both repelled at encoding and attracted at a later Bayesian integration stage (aligning with previously proposed models, Fritsche et al., 2020).

**Figure S4:**
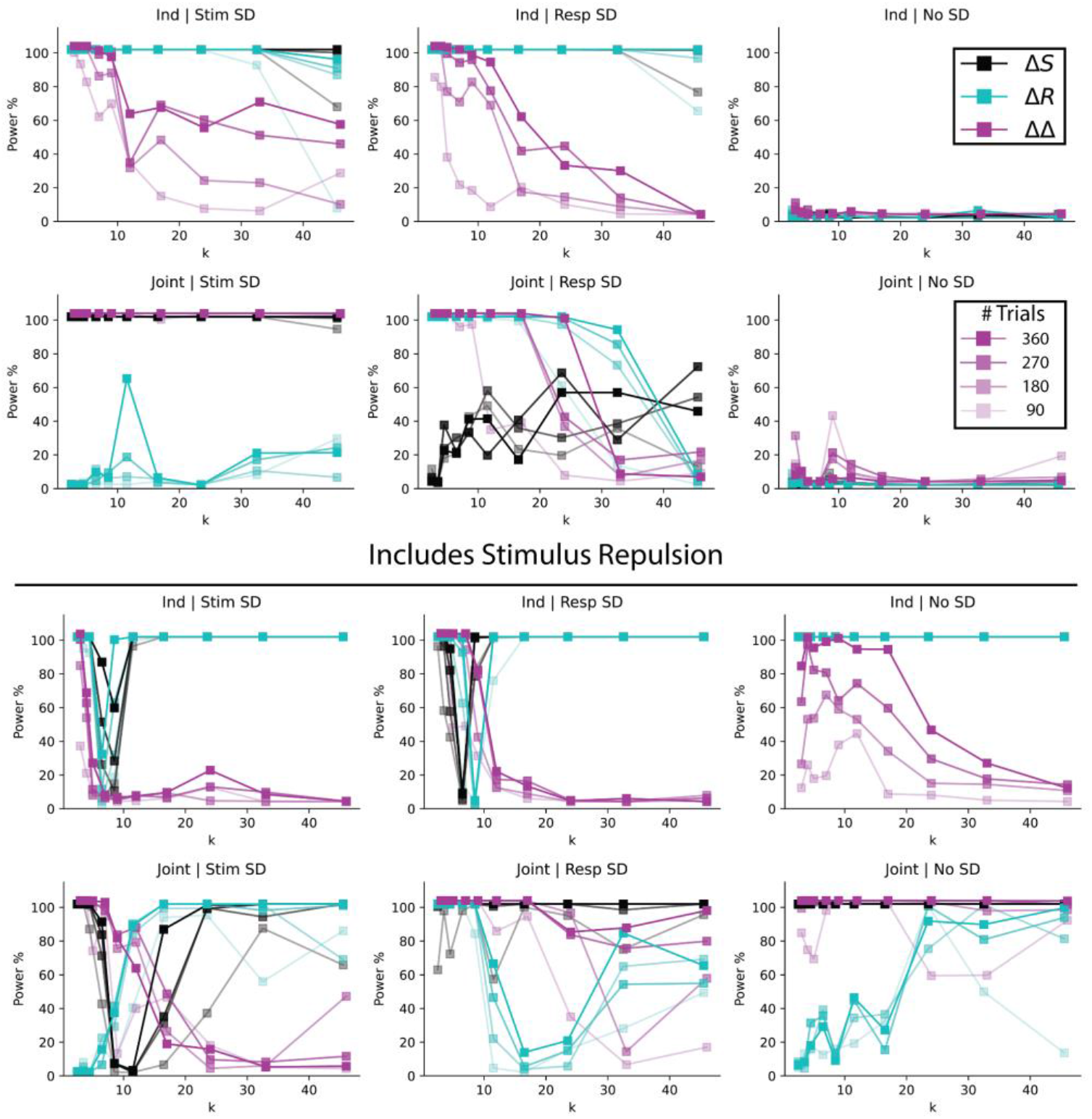
Expanded power analysis for observers without (top) and with (bottom) stimulus repulsion at encoding. Here we split out observers based on the number of trials completed per observer. Power values correspond to α=.001 for an experiment run with 30 participants.

**Figure S5:**
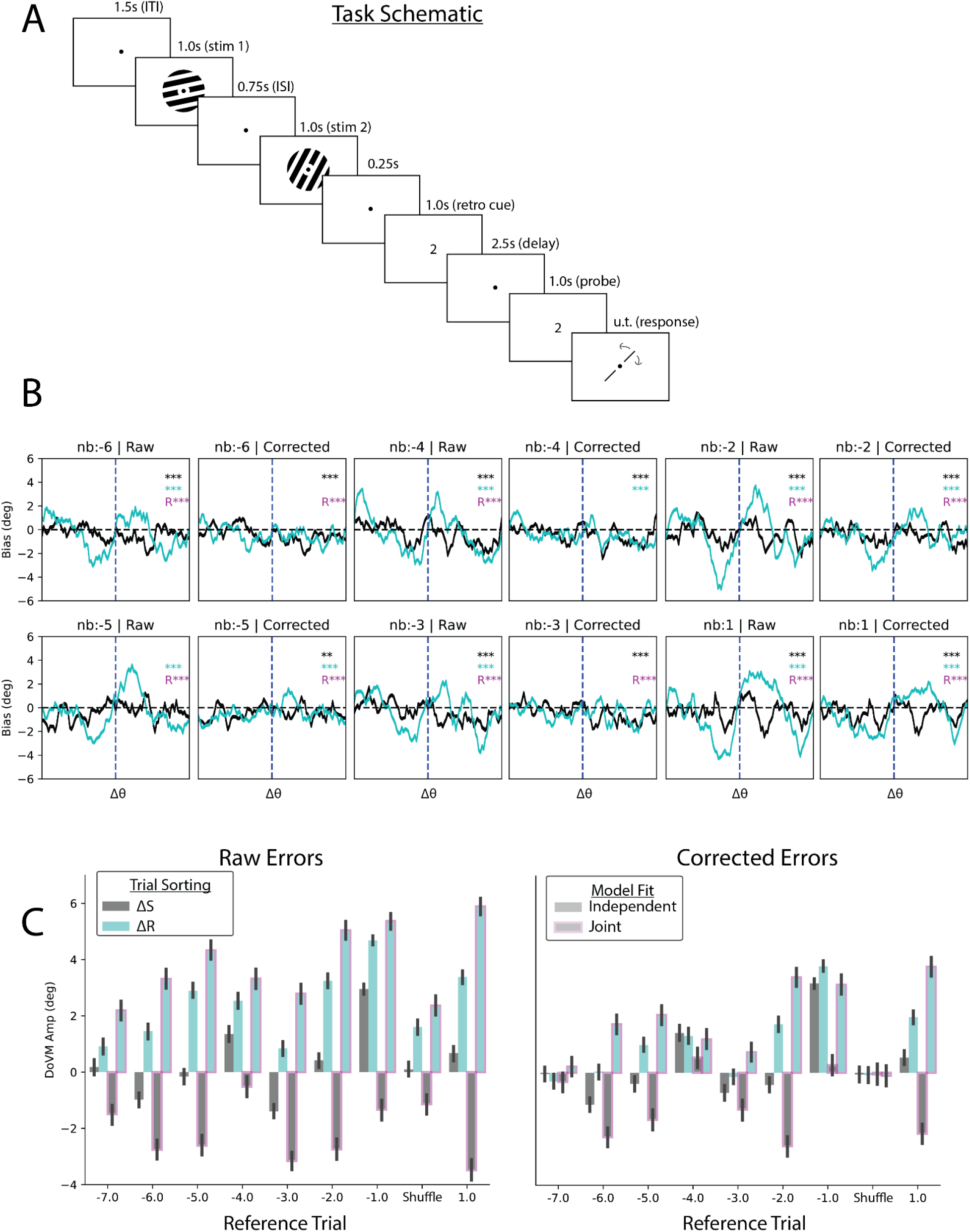
**A**. Full task schematic from delayed report paradigm. A Probabilistic retro-cue (80%) valid was presented immediately after the second item followed by a 100% valid probe and an untimed continuous report task controlled via the keyboard. Probe location initialized to a random location on each trial. **B**. Expanded stimulus and response bias curves for corrected and uncorrected errors for different number of trials back and using shuffled distribution. **C**. Quantified bias fits for both independent (no outline) and joint (magenta outline) models. Correcting errors removes spurious biases in the shuffled distribution (right, shuffle). Joint model reveals attraction to reported stimulus going back several trials.

## Footnotes

1 Note that here for simplicity we are equating the shape of the likelihood function, p(θ|m), with the posterior p(m|θ).

